# Structural basis for relief of the sarcoplasmic reticulum Ca^2+^-ATPase inhibition by phospholamban at saturating Ca^2+^ conditions

**DOI:** 10.1101/310607

**Authors:** Eli Fernández-de Gortari, L. Michel Espinoza-Fonseca

## Abstract

We have performed extensive atomistic molecular dynamics simulations to probe the structural mechanism for relief of sarcoplasmic reticulum Ca^2+^-ATPase (SERCA) inhibition by phospholamban (PLB) at saturating Ca^2+^ conditions. Reversal of SERCA-PLB inhibition by saturating Ca^2+^ operates as a physiological rheostat to reactivate SERCA function in the absence of PLB phosphorylation. Simulation of the inhibitory complex at super-physiological Ca^2+^ concentrations ([Ca^2+^]=10 mM) revealed that calcium ions interact primarily with SERCA and the lipid headgroups, but not with the cytosolic domain of PLB or the cytosolic side of the SERCA-PLB interface. At this [Ca^2+^], a single Ca^2+^ ion is translocated from the cytosol to the transmembrane transport sites. We used this Ca^2+^-bound complex as an initial structure to simulate the effects of saturating Ca^2+^ at physiological conditions ([Ca^2+^]_total≈_400 μM). At these conditions, ~30% of the Ca^2+^-bound complexes exhibit structural features that correspond to an inhibited state. However, in ~70% of the Ca^2+^-bound complexes, Ca^2+^ moves to transport site I, recruits Glu771 and Asp800, and disrupts key inhibitory contacts involving conserved PLB residue Asn34. Structural analysis showed that Ca^2+^ induces only local changes in interresidue inhibitory interactions, but does not induce dissociation, repositioning or changes in the structural dynamics of PLB. Upon relief of SERCA inhibition, Ca^2+^ binding produces a productive site I configuration that is sufficient for subsequent SERCA activation. We propose that at saturating [Ca^2+^] and in the absence of PLB phosphorylation, binding of a single Ca^2+^ ion in the transport sites rapidly shifts the equilibrium toward a non-inhibited SERCA-PLB complex.

## Introduction

The sarcoplasmic reticulum Ca^2+^-ATPase (SERCA) transports uses the energy derived from hydrolysis of one ATP to transport two Ca^2+^ ions from the cytosol into the lumen of the sarcoplasmic reticulum (SR) (1). In cardiac muscle cells, SERCA function is reversible regulated by the transmembrane phospholamban (PLB). PLB binds SERCA in a 1:1 heterodimeric regulatory complex and inhibits SERCA activity (2, 3). Phosphorylation of PLB then relieves SERCA inhibition (3) to increase the rate of cardiac muscle relaxation and to restore the SR Ca^2+^ load necessary for muscle contraction in subsequent beats (4).

PLB inhibits SERCA by binding to a large pocket located in the transmembrane (TM) domain of the pump (5–10). Spectroscopy studies have shown that in the bound complex, SERCA inhibition is tightly coupled to a structural transition between inhibitory (T) and non-inhibitory (R) structural states of PLB (11–14). More recently, x-ray crystallography studies showed that in its unphosphorylated form, PLB forms specific intermolecular interactions between conserved residue Asn34 and residue Gly801 of SERCA (15). Extensive studies by our group showed that these intermolecular interactions induce a substantial structural rearrangement of the transmembrane transport sites and stabilize a metal ion-free E1 intermediate of the pump protonated at residue Glu771, El·H^+^_771_ (16, 17). This SERCA intermediate serves as a kinetic trap that decreases SERCA’s apparent affinity for calcium at low Ca^2+^ concentrations and depresses the structural transitions necessary for Ca^2+^-dependent activation of SERCA (16, 17).

In the absence of PLB phosphorylation, relief of SERCA inhibition occurs at saturating Ca^2+^ concentrations (18, 19). Two mechanisms have been proposed for the relief of SERCA inhibition at saturating Ca^2+^ conditions: dissociation model and subunit model. The dissociation model proposes that PLB must dissociate from SERCA to relieve inhibition, whereas the subunit model hypothesizes that PLB acts as a functional subunit of SERCA, and inhibition is relieved by local structural rearrangements within the SERCA-PLB complex. Spectroscopic experiments in live cells, ER membranes, and reconstituted vesicles, support the subunit model, as they provide direct detection of the SERCA-PLB interaction at high Ca^2+^ concentrations (11,20–22). Crosslinking studies have suggested that saturation Ca^2+^ conditions induce PLB dissociation from SERCA, although they are also consistent with a structural rearrangement of the inhibitory complex (7, 19).

These studies indicate that relief of SERCA inhibition at saturating Ca^2+^ conditions requires structural changes that are difficult to determine with experiments alone. Therefore, in this study we designed a series of atomistic molecular simulations to determine the mechanisms for Ca^2+^-dependent relief of inhibitory interactions between SERCA and PLB. First, we used a full-length structure of SERCA bound to the inhibitory structure of unphosphorylated PLB as a starting structure to obtain a structure of the SERCA-PLB complex at saturating Ca^2+^ conditions. We then used the Ca^2+^-bound SERCA-PLB structural model generated through this computational approach to perform six independent 1-μs molecular dynamics (MD) simulations of the complex at physiologically relevant conditions. This set of independent simulations was used to systematically identify the effects of saturating Ca^2+^ conditions on the inhibitory contacts between SERCA and PLB in the absence of PLB phosphorylation.

## Results

### Interaction of Ca^2+^ with the SERCA-PLB complex at saturating Ca^2+^ concentrations

Physiologically relevant saturating free Ca^2+^ concentrations are in the low μM range (23), but these ion concentrations cannot be effectively modeled with our explicit MD simulations because the finite size of the systems would require less than one Ca^2+^ ion/system. To overcome this limitation, we first performed a 0.5-μs MD simulation of the SERCA-PLB complex at 10 mM CaCl2 to mimic saturating Ca^2+^ concentrations, and to match the experimental conditions previously used in x-ray crystallography studies (24, 25). At this Ca^2+^ concentrations, Ca^2+^ ions interact with several acidic residues of SERCA exposed on both sides of the lipid bilayer, as well as with the lipid headgroups (Figure 1). However, we found no indication that Ca^2+^ interacts with the cytosolic domain of PLB or near the cytosolic side of the SERCA-PLB interface (Figure 1).

**Figure 1.**
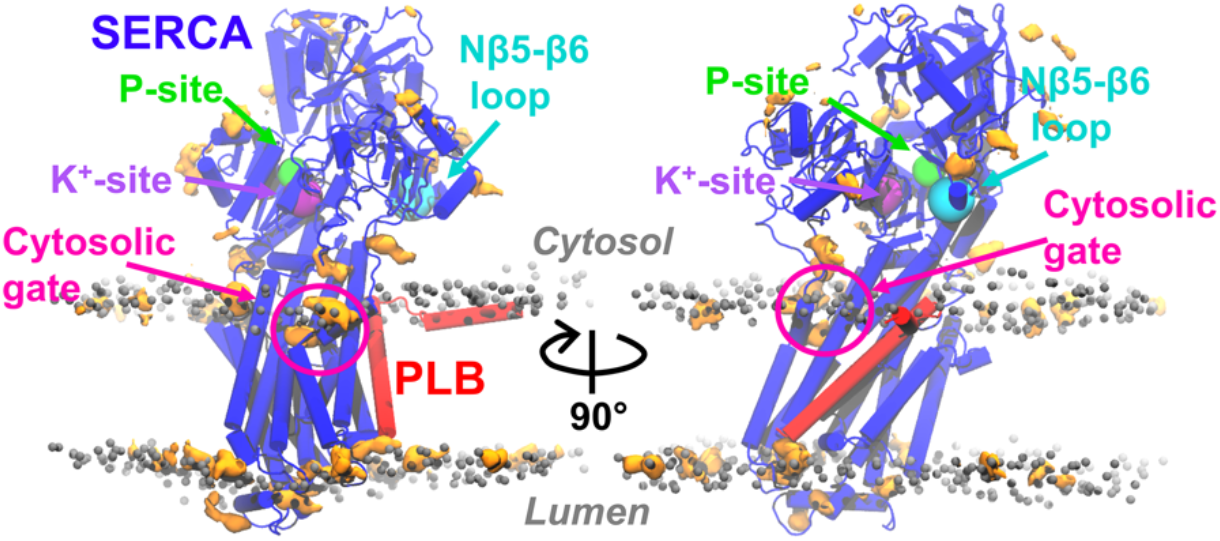
Map of the Ca^2+^-protein and Ca^2+^-lipid interactions in the MD trajectory of the complex at 10 mM Ca^2+^. The map of the weighted mass density of Ca^2+^ (orange surface representation), was calculated using a grid resolution of 1 Å and a cutoff distance of 3.5 Å between Ca^2+^ and the protein/lipid atoms. The headgroups of the lipid bilayer are shown as grey spheres; SERCA and PLB are shown in blue and red cartoons, respectively. The arrows indicate the location of functionally important sites in SERCA: phosphorylation site (Asp351) in green; the K+-binding site (Ala711, Ala714, Lys712 and Glu732) in purple; the Nβ5-β6 loop (Asp426–Lys436) in cyan; and the cytosolic gate (Asp59 and Glu309) in magenta.

Mapping of Ca^2+^-SERCA interactions revealed that Ca^2+^ ions do not form interactions with functionally important regions of the pump, such as the phosphorylation site (Asp351) or the K+-binding site involved in SERCA dephosphorylation (Ala711, Ala714, Lys712 and Glu732) (26) (Figure 1). In other cases, Ca^2+^ ions interact, albeit non-specifically, with other functional sites in the cytosolic headpiece such as the Nβ5-β6 loop (Asp426–Lys436) (27) (Figure 1). However, we found that Ca^2+^ ions bind to SERCA near the cytosolic gate that leads to the transport sites (Figure 1). On average, 2-3 Ca^2+^ ions occupy this region of the protein, but we found that only a single Ca^2+^ ion binds to residues Asp59 and Glu309 at the entrance of the cytosolic pathway (Figure 2A). We found that in the sub-μs timescale (t=0.34 μs), Glu309 translocates a single Ca^2+^ ion from Asp59 to Asp800 (Figure 2B). Ca^2+^ translocation is facilitated by a change in the dihedral angle N-Cα-Cβ-Cγ (χ1) of Glu309 from values of +165° and −165° to a χ1=-65°. Upon translocation, Ca^2+^ is stabilized in the transport sites by electrostatic interactions with residues Asp800 and Glu309 (Figure 2C). This Ca^2+^-bound configuration of the complex is stable for the remainder of the simulation time, so we use it as starting structure for six independent MD simulations, identified as CAL1 through CAL6. These MD simulations were used to determine the effects of saturating Ca^2+^ conditions at physiologically relevant conditions (e.g., ~400 μM Ca^2+^ and 100 mM K+).

**Figure 2.**
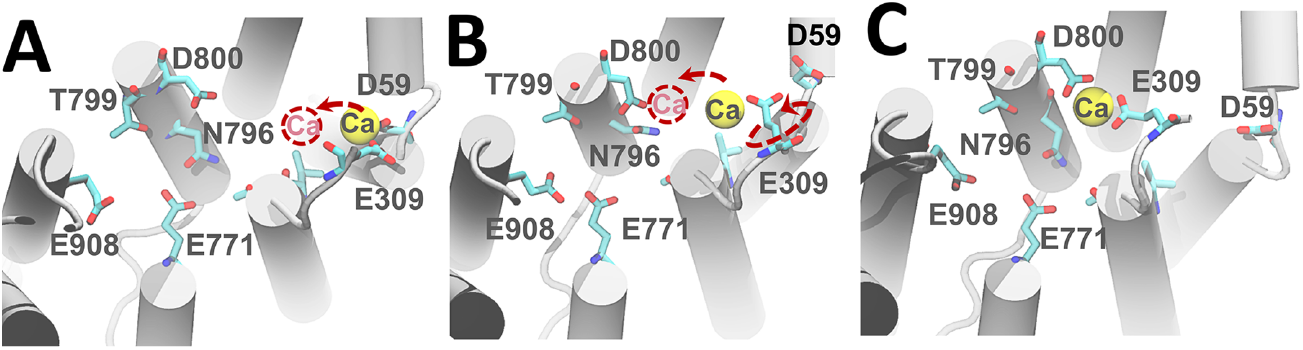
Mechanism for Ca^2+^ binding to the transport sites at super-physiological Ca^2+^ concentrations. (A) In the ns timescale, a single Ca^2+^ ion binds to SERCA residues Asn59 and Glu309, located at the entrance of the cytosolic pathway leading to the transport sites. (B) Following Ca^2+^ binding, rotation of the Glu309 sidechain facilitates translocation of Ca^2+^ through the cytosolic pathway into the transport sites. (C) Upon translocation, the position of Ca^2+^ in the transport sites is stabilized by electrostatic interactions with residues Asp800 and Glu309 for the remainder of the simulation time. In all panels, the TM helices are represented by grey ribbons, transport site residues are shown as sticks, and the Ca^2+^ ion is represented as a yellow sphere.

This set of MD simulations revealed that Ca^2+^ remains bound to the transport sites of SERCA and does not dissociate back to the cytosol at physiological conditions. We calculated the coordination number and coordination shell of Ca^2+^ to characterize the interactions that stabilize a single Ca^2+^ ion in the transport sites of SERCA. We define the coordination number of Ca^2+^ as the number of oxygen atoms within 3.5 Å of the calcium ion. This distance is normally considered to be the maximum possible distance between ligand oxygen atoms and the calcium ion (28–30). We found that in all trajectories, Ca^2+^ interacts with the transport sites predominantly with a coordination number of 7, although Ca^2+^ also exhibits a coordination number of 8 in a small percentage of the total simulation time (<10%). These coordination numbers fall within the typical values estimated from Ca^2+^-protein (31, 32) and Ca^2+^-SERCA complexes (24, 33).

In two trajectories, CAL1 and CAL3, the Ca^2+^ ion interacts predominantly with seven coordinating oxygen atoms primarily in a pentagonal bipyramidal coordination geometry, in agreement with previous crystallographic studies of Ca^2+^-bound **SERCA** (33) (Figure 3A). The seven coordinating ligands for Ca^2+^ are the carboxylic oxygen atoms from residues Glu309 and Asp800, the carbonyl moiety from residue Asn796, and between 2 and 4 water molecules (Figure 3A). We found that in trajectories CAL2, and CAL4 through CAL6, Ca^2+^ also coordinates to oxygen atoms within the transport site predominantly in a heptavalent pentagonal bipyramidal geometry. However, in these four trajectories, Glu771 rapidly (t=25-250 ns) replaces Glu309 in the first coordination shell of Ca^2+^ (Figure 3B). In this binding mode, Ca^2+^ interacts with the oxygen atoms from transport site residues Glu771 and Asp800, the carbonyl group from residue Asn796, and 2-3 water molecules (Figure 3B).

**Figure 3.**
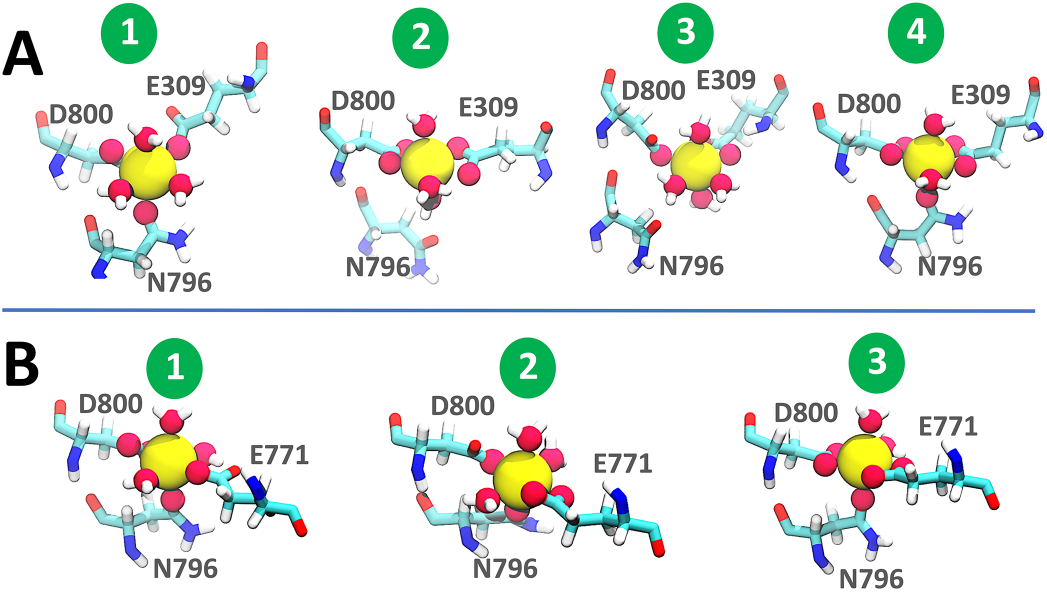
Structure and ligand coordination of Ca^2+^ in the transport sites of PLB-bound SERCA simulated at physiological conditions. (A) Representative interactions of Ca^2+^ with residues Glu309, Asn796 and Asp800, as well as water molecules in the transport sites. (B) Structure of the most populated Ca^2+^ coordination geometries involving SERCA residues Glu771, Asn796 and Asp800, and 2-3 water molecules. In all panels, transport site residues and water molecules are shown as sticks, oxygen atoms in the first coordinating shell of Ca^2+^ are shown as pink spheres, and the Ca^2+^ ion is represented as a yellow sphere.

### Effect of saturating Ca^2+^ conditions on PLB-induced inhibitory interactions

Recent studies have shown that PLB residue Asn34, which is absolutely required for SERCA inhibition (34), forms specific hydrogen bond interactions with Gly801 and Thr805 in the TM domain of SERCA (15–17). These interactions induce alterations in the transport site geometry that prevent metal ion occlusion in the transport sites (16, 17). The SERCA-Ca^2+^ interactions shown in Figure 3 suggest that saturating Ca^2+^ concentrations alter SERCA-PLB inhibitory contacts at physiological conditions. Therefore, we measured distances between intermolecular residue pairs Gly801-Asn34 and Thr805-Asn34. SERCA residues Glu771 and Asp800 play a key role in Ca^2+^ occlusion in the transport sites (35), so we also measured the interresidue distance Glu771-Asp800.

In those cases where Ca^2+^ primarily binds to SERCA residues Glu309 and Asp800 (trajectories CAL1 and CAL3), PLB residue Asn34 interacts directly with the backbone oxygen of Gly801 and the sidechain hydroxyl group of Thr805 (Figure 4). In both cases, the intermolecular interactions Asn34-Gly801 and Asn34-Thr805 are present for most of the simulation time. Furthermore, we found that the carboxyl groups of transport site residues Glu771 and Asp800 in these trajectories are separated by a distance of at least 9 Å (Figure 4). This spatial separation between Glu771 and Asp800 which is characteristic of the inhibited SERCA-PLB complex in the absence of Ca^2+^ (15, 16). Therefore, the stability of the inhibitory interactions and the large spatial separation between Glu771 and Asp800 indicate that the structures populated in the trajectories CAL1 and CAL3 correspond to those of an inhibited Ca^2+^-bound state of the SERCA-PLB complex (15–17).

**Figure 4.**
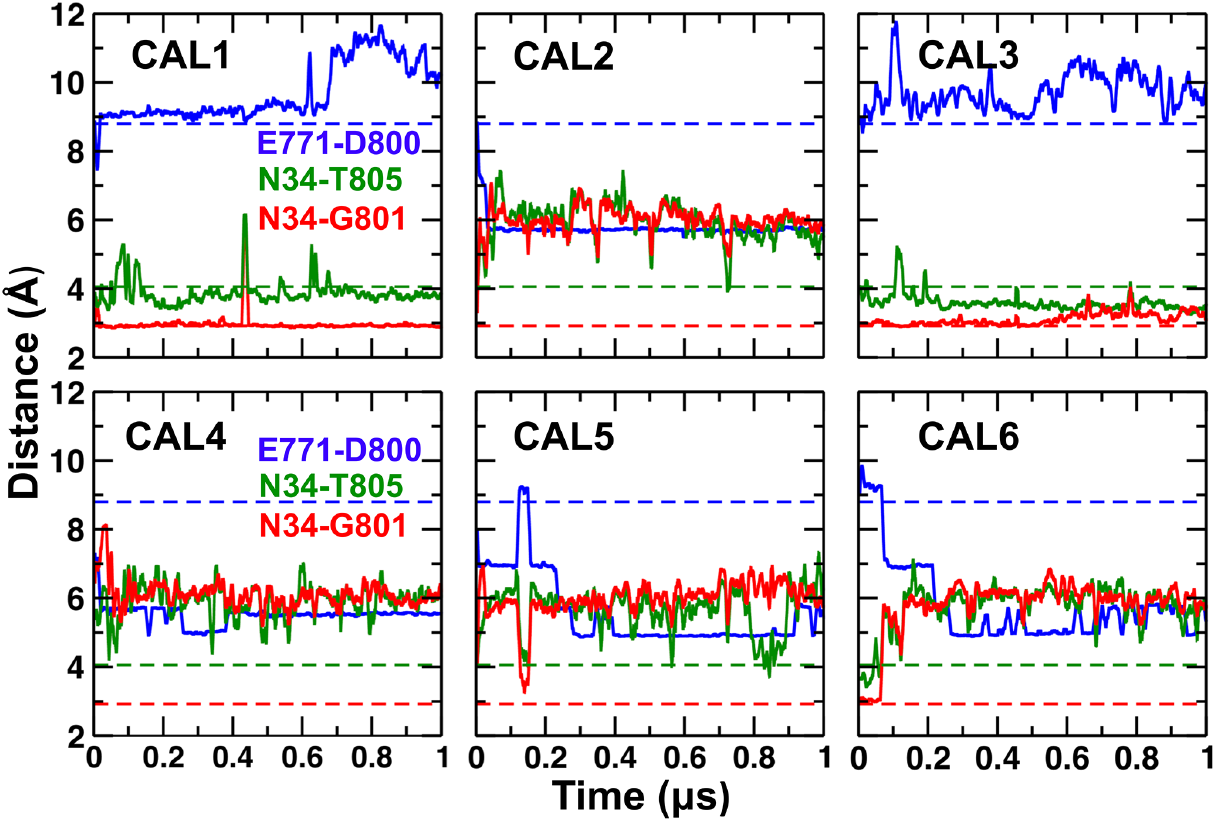
Distance evolution between residue pairs associated with inhibition Ca^2+^ occlusion calculated from the MD trajectories of the Ca^2+^-bound SERCA-PLB complex. Distances between key inhibitory contacts involving conserved PLB residue Asn34 and SERCA Gly801 were calculated using atoms Nδ of Asn34 and the backbone oxygen of Gly801. The distance between Thr805 and Asn34 was calculated between the Oγ of Thr805 and Nδ of Asn34. Distances between Glu771 and Asp800, which occlude Ca^2+^ in site I, were calculated using atoms Cδ and Cγ, respectively. The dashed lines indicate the interresidue distances in the initial structure inhibited complex reported in a previous study by our group (16).

In four MD trajectories, CAL2, CAL4, CAL5 and CAL6, we found substantial changes in the distances between intermolecular residue pairs Asn34-Gly801 and Asn34-Thr805. Here, the distance Asn34-Thr805 increases by 1-2 Å (Figure 4); however, the most significant change is the 3-Å increase in the distance between the sidechain of Asn34 and the backbone oxygen of Gly801 (Figure 4). Most importantly, we found that the spatial separation Asn34 of PLB and Gly801 of SERCA in these trajectories occurs concomitantly with a 3-4 Å decrease in the distance between transport site residues Glu771 and Asp800 (Figure 4). These changes in interresidue distances are also accompanied by large shift in the dihedral angle Cα-Cβ-Cγ-Nδ (χ2) of Asn34 from two narrow distributions at χ2=+180° and χ2=-180° to a single broad distribution with mean around −20° (Figure 5). This structural change is mostly characterized by a transition from an extended side chain conformation to a self-contact (36) involving side-chain nitrogen and backbone oxygen atoms of Asn34 (Figure 5). These findings indicate that in trajectories CAL2, CAL4, CAL5 and CAL6, the sidechain of PLB residue Asn34 becomes more mobile and no longer establishes inhibitory contacts with SERCA.

**Figure 5.**
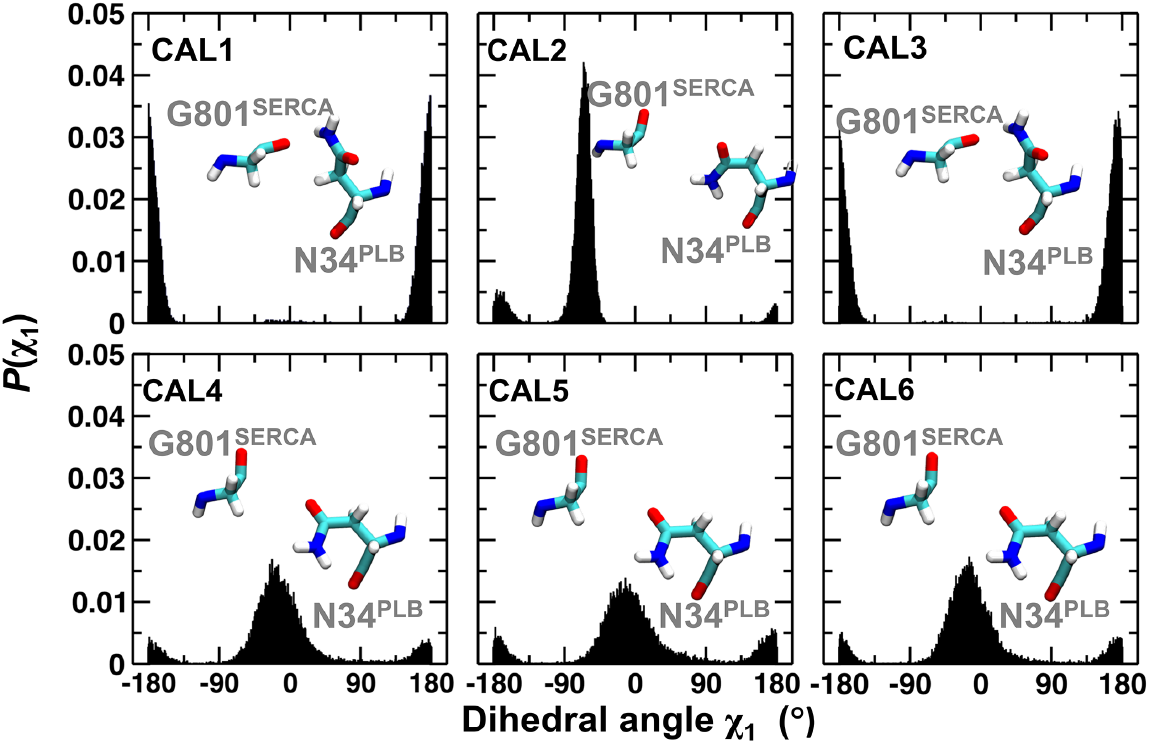
χ_2_ dihedral angle probability distributions for PLB residue Asn34 in the trajectories of the Ca^2+^-bound SERCA-PLB complex. The insets illustrate both the preferred conformation adopted by Asn34 and the formation or disruption of the inhibitory interaction Asn34-Gly801 in each individual MD trajectory.

Taken together, these observations provide evidence which is consistent with relief of SERCA-PLB inhibition at saturating Ca^2+^ conditions. However, this phenomenon is not consistently observed in all six trajectories, which suggests that this disruption of inhibitory contacts probably occurs in equilibrium and in a Ca^2+^-independent manner. To test this hypothesis, we performed six 1-μs MD simulations of the SERCA-PLB complex in the absence of Ca^2+^. We found that while intermolecular distance Asn34-Thr805 varies between independent MD trajectories, Asn34 of PLB consistently remains spatially close to SERCA residue Gly801 throughout the entire simulation time in all trajectories (Figure 6). We also found that carboxylic groups of transport site residues Glu771 and Asp800 is are separated by a distance larger than 9 Å (Figure 6); such interresidue distances are longer than the 6-Å separation required for Ca^2+^ occlusion (24, 33). These findings indicate that relief of SERCA-PLB inhibitory contacts does not occur spontaneously in the absence of Ca^2+^, and that Ca^2+^ binding to SERCA at saturating Ca^2+^ conditions disrupts key SERCA-PLB inhibitory contacts at physiological conditions.

**Figure 6.**
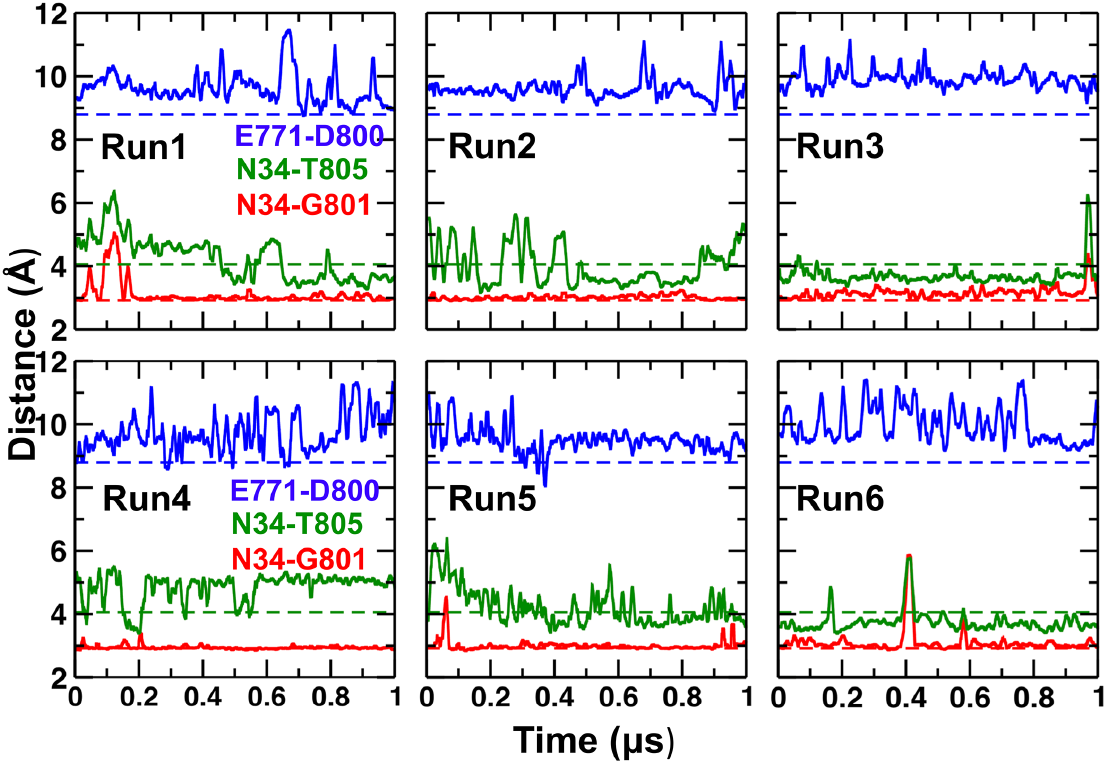
Distance evolution between residue pairs associated with inhibition Ca^2+^ occlusion calculated from the MD trajectories of the Ca^2+^-free SERCA-PLB complex. Interresidue distances were calculated using the same atom pairs described in **Figure 4**. The dashed lines indicate the distances in the initial structure inhibited complex reported in reference (16). The plots show that in the absence of Ca^2+^, the inhibitory contacts between Asn34 of PLB and Gly801 are not disrupted throughout the entire simulation time in all trajectories.

### Effects of saturating Ca^2+^ conditions on the structural dynamics of PLB

Our results demonstrate that Ca^2+^ binding to the transport sites of SERCA at saturating Ca^2+^ conditions generally disrupt key inhibitory interactions between SERCA and PLB. However, it is not clear if disruption of the inhibitory interactions is linked to (i) changes in the structural dynamics of PLB, (ii) a reorganization of the SERCA-PLB interface (e.g., PLB dissociation), or (iii) local changes involving interresidue interactions along the interface. Therefore, we performed extensive measurements of structural parameters to determine the changes in the structural dynamics of PLB in response to saturating Ca^2+^ conditions.

In the both inhibited and non-inhibited Ca^2+^-bound complexes, the cytosolic and TM helices that contain the regulatory phosphorylation and inhibitory domains populate an α-helical structure for >95% of the time. Average interhelical angles between the cytosolic (Val4-Thr17) and TM (residues Arg25–Leu52) helices of PLB fluctuate between 52° and 77° (Table 1), which correspond to a T-shaped architecture of PLB (Figure 7A). We found that the calculated interhelical angles of PLB in both inhibited and non-inhibited complexes are within the range of those determined experimentally for the unphosphorylated PLB monomer in solution (37, 38). These findings are in agreement with previous spectroscopic studies (11, 39), and demonstrate that saturating Ca^2+^ conditions do not induce order-to-disorder transitions associated with PLB phosphorylation (12,39–43).

**Figure 7.**
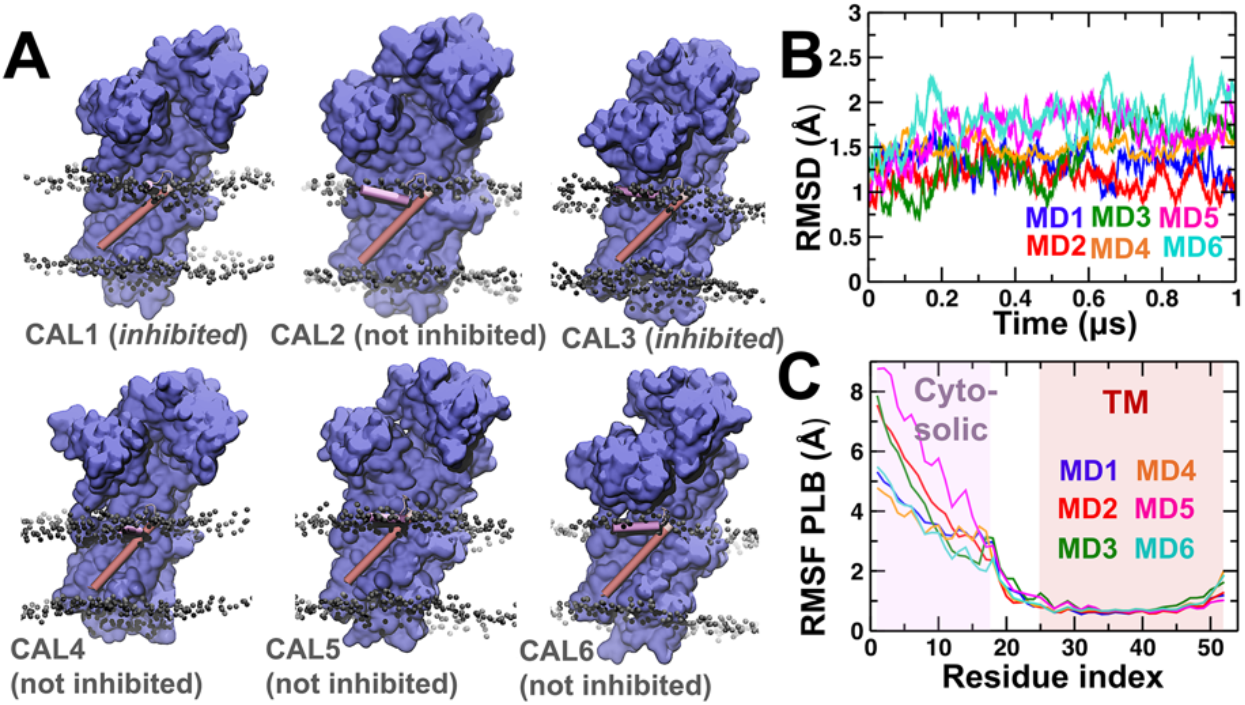
Effects of saturating Ca^2+^ conditions on the structural dynamics of PLB at physiologically relevant conditions. (A) Structures of the Ca^2+^-bound SERCA-PLB complex embedded in a lipid bilayer at the end of the each 1-μs MD trajectory. The headgroups of the lipid bilayer are shown as grey spheres; SERCA is shown as a blue surface representation. PLB is shown as a cartoon representation and colored according to its functional domains: cytosolic (pink) and TM (red). Based on the structural analysis reported in this study, each structure is labeled as inhibited or not inhibited. (B) Time-dependent RMSD evolution of the TM helix of PLB in the Ca^2+^-bound SERCA-PLB complexes. The RMSD was calculated using backbone alignment for TM helices of the SERCA. (C) Ca RMSF values of PLB calculated from each independent MD trajectory. The shaded areas show the location of the cytosolic and TM helices of PLB.

**Table 1.** Effects of saturating Ca^2+^ conditions on the structural dynamics of PLB and the inhibitory SERCA-PLB contacts

| Trajectory | PLB<br>interhelical<br>angle (°)<sup>a,b</sup> | PLB tilt<br>angle (°)<sup>a,c</sup> | Fraction of native inhibitory contacts, $Q_{inh}^{a,d}$ | | | |
| --- | --- | --- | --- | --- | --- | --- |
|  |  |  | Leu31 | Asn34 | Phe35 | Ile38 |
| CAL1 | 77.2 ± 8.1 | 4.5 ± 1.2 | 0.85±0.06 | 0.84±0.05 | 0.86±0.10 | 0.87±0.05 |
| CAL2 | 60.0 ± 8.5 | 3.1 ± 1.2 | 0.73±0.09 | 0.55±0.07 | 0.68±0.10 | 0.79±0.06 |
| CAL3 | 51.5 ± 10.5 | 5.4 ± 2.1 | 0.81±0.08 | 0.86±0.06 | 0.89±0.07 | 0.84±0.06 |
| CAL4 | 72.0 ± 7.5 | 2.5 ± 1.2 | 0.71±0.09 | 0.52±0.06 | 0.71±0.12 | 0.73±0.09 |
| CAL5 | 55.1 ± 9.3 | 3.9 ± 1.5 | 0.70±0.13 | 0.62±0.06 | 0.79±0.06 | 0.82±0.06 |
| CAL6 | 56.5 ± 8.7 | 2.3 ± 1.2 | 0.73±0.11 | 0.54±0.05 | 0.70±0.07 | 0.73±0.09 |

<sup>a</sup> Values reported as average ± standard deviation.
<sup>b</sup> Interhelical angle is defined as the polar angle between the cytosolic (Val4-Thr17) and TM (residues Arg25–Leu52) helices of PLB.
<sup>c</sup> Axial tilt angle was calculated using backbone alignment, with SERCA–PLB crystal structure [PDB: 4kyt, (15)] as a reference.
<sup>d</sup> These PLB residues play important roles in inhibition of SERCA (45)

We measured time-dependent root-mean square deviation (RMSD) to determine the extent to which the position of the TM domain of PLB changes in the trajectories of the Ca^2+^-bound complexes. RMSD plots revealed that the position of PLB in the binding groove in all six trajectories does not deviate substantially (RMSD<2.5 Å) from that determined by x-ray crystallography (Figure 7B). In addition, root-mean square fluctuation (RMSF) calculations using the main-chain Cα atoms show that the cytosolic helix of PLB is highly mobile in solution (Figure 7C). This behavior corresponds to the inherent diffusion of the helical domain through the viscous bilayer surface, and it is uncorrelated with the presence or absence of inhibitory contacts. The RMSF values of the TM domain residues in all MD trajectories are substantially smaller than those calculated for the cytosolic helix; this indicates that the TM domain of PLB has very low mobility in the μs time scale. The RMSF values calculated for the TM domain of PLB, and particularly those around the residue Asn34, are virtually identical both in the presence and absence of intermolecular inhibitory contacts (Figure 7C).

We also calculated changes in the tilt angle of the TM helix of PLB to complement RMSD and RMSF measurements. We measured he relative tilt angle of the TM helix using the crystal structure of SERCA-PLB as a reference (Table 1). We found that the TM helix exhibits on average a 3.6° increase in the tilt angle relative to the position of PLB in the crystal structure of the complex (Table 1). There is no correlation between the loss of inhibitory contacts and the small change in tilt angle, which indicates that saturating Ca^2+^ conditions do not have an effect on the position of PLB in the complex. Therefore, it is likely that the small changes in tilt angle correspond to the effects of annular lipid shell on the structural dynamics of the SERCA-PLB interface (44).

Relief of inhibitory contacts is not accompanied by large changes in the structure of PLB in the complex, so disruption of SERCA-PLB inhibitory contacts must occur locally at the interface of the complex. To test this hypothesis, we measured the fraction of native inhibitory contacts, *Qinh*, between SERCA and PLB residues Leu31, Asn34, Phe35 and Ile38; these residues, located near the cytosolic side of the complex, play important roles in inhibition of SERCA (45). Analysis of the *Qinh* values showed that in the complexes with intact inhibitory contacts (CAL1 and CAL3), there is a high retention of native inhibitory contacts *(Qinh* >0.8) between key PLB residues and SERCA (Table 1). As anticipated, there is a substantial decrease in native inhibitory contacts (Q_inh_=0.52-0.62) for PLB residue Asn34 in the trajectories where inhibitory contacts are disrupted (CAL2, CAL4, CAL5 and CAL6, Table 1). However, we found that a loss in native inhibitory contacts also occurs, albeit more moderately, at PLB positions Leu31, Phe35 and Ile38 (Table 1). This indicates that saturating Ca^2+^ conditions primarily affect local intermolecular interactions involving the side chain of PLB residue Asn34, but also affect intermolecular interactions involving PLB residues within the inhibitory site at the SERCA-PLB interface.

### Structure of the transport sites upon Ca^2+^-induced disruption of SERCA-PLB inhibitory interactions

We determined if the structural changes following disruption of inhibitory SERCA-PLB interactions correspond to those associated with the formation of competent transport sites. To this aim, we measured the RMSD values for residues in sites I and II between the MD trajectories and the crystal structure of SERCA bound to two Ca^2+^ ions (E1·2Ca^2+^, PDB: 1su4). We also performed distance measurements to determine if the location of the Ca^2+^ in the MD trajectories correspond to that determined by x-ray crystallography (33).

Visualization of the transport sites in the trajectories CAL1 and CAL3, where the inhibitory SERCA-PLB interactions are intact, show that sites I and II are collapsed (Figure 8). Calculated RMSD values for sites I and II (RMSD>2.7 Å) indicate that there is a poor overlap between these trajectories and the crystal structure of E1·2Ca^2+^ (Table 2). In these trajectories, the position of Ca^2+^ does not overlap with any of the two sites resolved by x-ray crystallography (Figure 8). Instead, the Ca^2+^ ion binds to a location that is distant from sites I (r≈5.5 Å) and site II (r≈3.5 Å). These measurements indicate that in the presence of inhibitory SERCA-PLB contacts, Ca^2+^ does not bind to either site I or II, and the residues in the transport sites adopt a non-competent structure.

**Figure 8.**
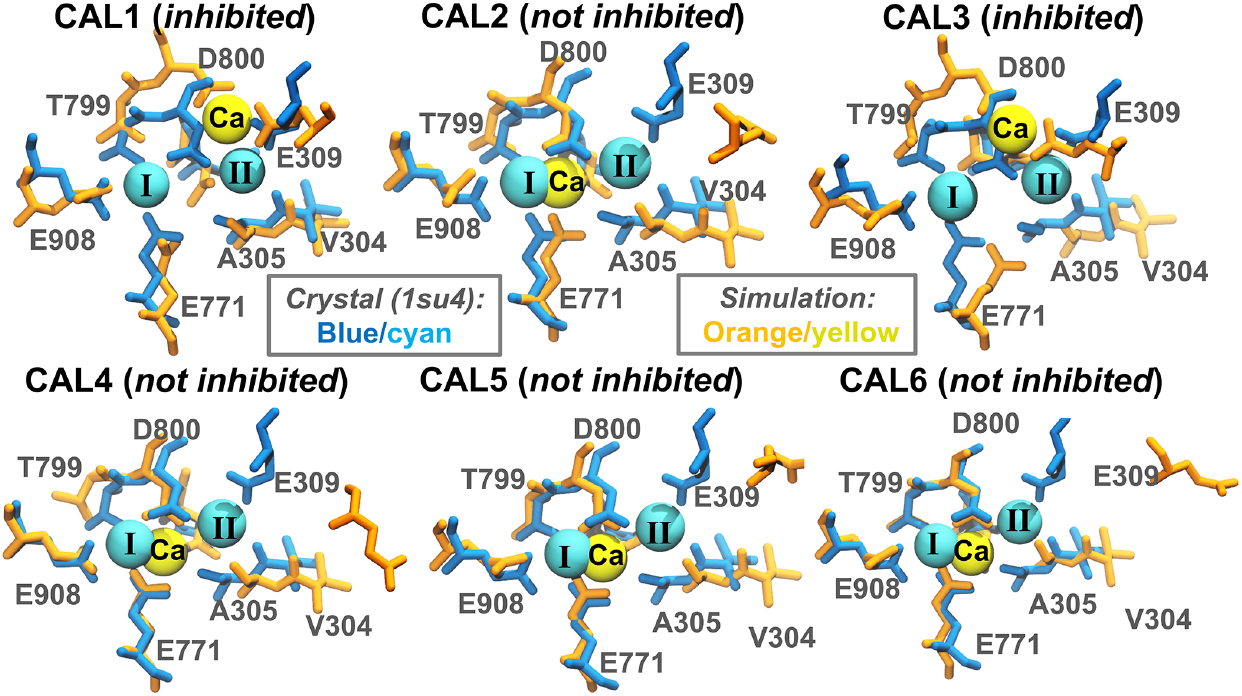
Ca^2+^ interactions with the transport sites of SERCA in the SERCA-PLB complex. Representative structures extracted from each MD simulation illustrate the location of Ca^2+^ (yellow sphere) and the structural arrangement of the residues that occlude Ca^2+^ in the transport sites (orange sticks). For comparison, each MD structure was superposed on the crystal structure of the Ca^2+^-bound SERCA, E1·2Ca^2+^ (33), to show the structure and location of the competent transport sites I and II; the side chains and Ca^2+^ ions resolved in the crystal structure are shown as blue sticks and cyan spheres, respectively. Based on the structural characteristics of each configuration, the structures of the transport sites are labeled as inhibited or not inhibited.

**Table 2.** Structural features of the transport sites at saturating Ca^2+^ conditions

| Trajectory | RMSD of heavy atoms<br>(Å)<sup>a,b</sup> | | Distance, $r$ (Å)<sup>a,c</sup> | |
| --- | --- | --- | --- | --- |
|  | Site I | Site II | Ca<sup>2+</sup>-site I | Ca<sup>2+</sup>-Site II |
| CAL1 | 2.8 ± 0.2 | 2.7 ± 0.3 | 5.5 ± 0.9 | 3.1 ± 0.5 |
| CAL2 | 2.2 ± 0.3 | 4.4 ± 0.5 | 1.9 ± 0.2 | 4.2 ± 0.2 |
| CAL3 | 3.2 ± 0.1 | 4.6 ± 0.2 | 5.6 ± 0.6 | 4.0 ± 0.4 |
| CAL4 | 2.1 ± 0.1 | 4.7 ± 0.3 | 2.2 ± 0.2 | 4.2 ± 0.3 |
| CAL5 | 1.7 ± 0.1 | 3.8 ± 0.3 | 2.2 ± 0.3 | 3.8 ± 0.3 |
| CAL6 | 2.0 ± 0.1 | 4.9 ± 0.5 | 2.1 ± 0.3 | 4.0 ± 0.2 |

<sup>a</sup> Values reported as average ± standard deviation.
<sup>b</sup> RMSD was calculated using heavy atom alignment, with E1·2Ca<sup>2+</sup> crystal structure [PDB: 1su4 (33)] as reference.
<sup>c</sup> Distances relative to the position of Ca<sup>2+</sup> in sites I and II determined by x-ray crystallography (33).

In the MD simulations where the inhibitory SERCA-PLB are disrupted (trajectories CAL2, CAL4, CAL5 and CAL6), the structure of transport site II also deviates substantially from that in the crystal structure of E1·2Ca^2+^ (Figure 8 and Table 2). However, we found that Ca^2+^ ion binds in a reproducible manner to a location that partially overlaps that of site I (r≈2 Å) determined by x-ray crystallography (Figure 8 and Table 2). When a single Ca^2+^ ion binds to this site, residues Glu771, Thr799, Asp800 and Glu908 of site I adopt a structural arrangement that is similar to that found in the crystal structure of the PLB-free E1·2Ca^2+^ state of SERCA (RMSD<2.2 Å, Figure 8). The structural rearrangements in the transport sites that follow Ca^2+^-induced relief of SERCA inhibition are reproducible in trajectories CAL2, and CAL4 through CAL6 (Figure 8), and correspond to the formation of a competent site I.

## Discussion

We present a mechanistic study of the SERCA-PLB regulatory interactions at saturating Ca^2+^ conditions, thus providing quantitative insight into fundamental processes of activation of Ca^2+^ transport in the heart. We show that in a solution containing 10 mM Ca^2+^, calcium ions interact primarily with both cytosolic and luminal sides of SERCA and the lipid headgroups. Here we show for the first time (to our knowledge) that at super-physiological Ca^2+^ conditions, Ca^2+^ ions interact with the luminal C-terminal region of PLB, but not with the cytosolic domain of PLB or the cytosolic side of the SERCA-PLB interface. This indicates that Ca^2+^ does not compete with PLB at the interface of the complex and does not have a direct effect on the structural dynamics and stability of unphosphorylated PLB. Previous FRET spectroscopy experiments support our data and show that saturating Ca^2+^ conditions do not alter the structural dynamics of either unphosphorylated PLB or the stability of the SERCA-PLB heterodimer (11).

At [Ca^2+^]=10 mM, Ca^2+^ ions show a strong preference for binding to SERCA residues Asp59 and Glu309, located near the entrance of the pathway that connects the cytosol with the transmembrane transport sites (46). The affinity of these residues for Ca^2+^ is supported by mutagenesis studies showing that Asp59 plays a central role in recruiting Ca^2+^ ions (47). We found that Glu309 translocates a single Ca^2+^ ion from Asp59 to Asp800, a critical residue in the transport sites that coordinates Ca^2+^ at sites I and II (33). This mechanism for Ca^2+^ translocation is in qualitative agreement with Brownian dynamics studies showing that fast Ca^2+^ binding to the transport sites is primarily guided by Glu309 (48).

In the absence of adequate charge neutralization of the transmembrane transport sites, SERCA denaturalization occurs very rapidly even within the native membrane at physiological pH (49, 50). Previous studies have shown that this electric charge can be compensated in the absence of Ca^2+^ by transport site protonation (16, 17), or by binding of metal ions K+ (51), Na+ (52–54) or Mg^2^+ (55, 56). This suggests that super-physiological concentrations of Ca^2+^ used in this study simply satisfy transport site charge neutralization, and that the Ca^2+^-bound state of the SERCA-PLB complex might not represent a functional state in the cell. However, we found that even at such high Ca^2+^ concentrations, only a single Ca^2+^ ion occupies the transport sites at a time. This finding is consistent with previous studies showing that the Ca^2+^ binding to SERCA occurs in a sequential manner (57–59), and indicates that the Ca^2+^-bound SERCA-PLB structure represents a functional state of the complex at saturating Ca^2+^ conditions. In the absence of other Ca^2+^ ions, a single Ca^2+^ bound to the SERCA-PLB complex produces to a total |Ca^2+^| of ~400 μM. This value falls in the middle of previous estimates at elevated cytosolic Ca^2+^ in the cardiomyocyte (60–62), so we used the Ca^2+^-bound SERCA-PLB complex as a starting structure to probe the structural mechanism for relief of SERCA inhibition by PLB.

We found that in 30% of the MD trajectories, Ca^2+^ binds to Glu309 and Asp800 in an orientation that is similar to that original found at Ca^2+^ concentrations of 10 mM. In this configuration, the SERCA-PLB inhibitory contacts remain intact in the μs time scale, and transport sites I and II lack the competent structural organization that is distinctive of the Ca^2+^-bound state of SERCA (33,54,63). This structural arrangement of the corresponds to an Ca^2+^-bound, inhibited structure of the SERCA-PLB complex. These findings provide new insights into the mechanism for PLB inhibition of Ca^2+^-dependent activation of SERCA at physiological conditions. For instance, previous experimental studies suggesting that PLB suppresses Ca^2+^ binding to SERCA (64), yet PLB binding decreases SERCA’s apparent affinity for Ca^2+^ only by 2-to 3-fold in the μM range (65). Our simulations help reconcile these models, since it shows that PLB-induced changes in the transport sites delay Ca^2+^ binding to either sites I or II, thus altering the apparent Ca^2+^ affinity of SERCA (34).

We found that the SERCA-PLB inhibitory interactions are disrupted in 70% of the MD trajectories of the Ca^2+^-bound complex. In these cases, the initially bound Ca^2+^ moves to site I and recruits the carboxylic groups of transport site residues Glu771 and Asp800. These Ca^2+^-induced structural changes occur concomitantly with a loss in the intermolecular interaction between the side chain of PLB residue Asn34 and the backbone oxygen of SERCA residue Gly801. Contrary to previous studies, our data indicates that this Ca^2+^-dependent relief inhibitory contacts does not result from PLB dissociation (19), large structural rearrangements in the SERCA-PLB interface (22), or changes in the native structural dynamics of PLB in the complex. Instead, PLB remains bound to SERCA, but PLB residue Asn34 becomes dynamically more disordered and is unable to establish inhibitory contacts with SERCA. FRET spectroscopy experiments support these findings, and show that Ca^2+^ acts upon SERCA-PLB complex exclusively at the TM domain level, that PLB does not dissociate from SERCA, and that Ca^2+^ does not induce phosphorylation-like order-to-disorder transitions in the cytosolic domain of PLB (11). These structural changes occur at much smaller spatial scale than that accessible by spectroscopy, so we propose that the Ca^2+^-induced repositioning and mobility of PLB residue Asn34 observed in our simulations can be verified experimentally by x-ray crystallography.

What are the specific interactions between Ca^2+^ and the transport sites that induce relief of inhibition? The crystal structure of the complex between sarcolipin (SLN), a PLB analog, and SERCA revealed a single Mg^2+^ ion bound to transport site residues Glu771 and Asp800 (56). In this structure, the intermolecular inhibitory interactions are partially altered, which suggests that binding of divalent metal ions in the transport sites is sufficient to reverse SERCA inhibition. However, extensive studies by our group have demonstrated that in the inhibitory complex, Mg^2+^ adopts a rigid octahedral coordination geometry that has a preference for binding water molecules as opposed to bulky protein side chain dipoles (17). In addition, the ionic radius of Mg^2^+ is smaller than that of Ca^2+^ (66), so adding side chain dipoles to the coordination shell is thermodynamically more favorable for Ca^2+^ than for Mg^2+^, so Ca^2+^ can produce drier, bulkier coordination complexes (67, 68). This explains why Ca^2+^, but not Mg^2^+, recruits both Glu771 and Asp800 in the transport sites (17, 55). Owing to these distinctive properties of Ca^2+^, the tug-of-war between the attraction of Glu771 and Asp800 for Ca^2+^ drag Gly801 backbone along with them as they move in toward Ca^2+^. These Ca^2+^-induced changes destabilize the interaction between Gly801 and PLB residue Asn34 and induce relief of SERCA inhibition by PLB. It is our postulate that Glu771 and Asp800 to a large degree define the range of coordination spheres that help preserve or disrupt the inhibitory contacts in the SERCA-PLB complex.

Finally, we ask whether the mechanism for relief of SERCA inhibition detected in our simulations produce an intermediate state in pathway toward SERCA activation. First, we found that upon relief of inhibitory contacts, the sidechain of Glu309 populates a geometry in which the carboxylic group points toward the cytosol. We propose that this orientation of Glu309 is essential for binding and gating of a second Ca^2+^ ion in the transport site II (69). Second, we found that relief of inhibitory SERCA-PLB interactions occurs only when a single Ca^2+^ binds near transport site I, and in agreement with mutagenesis studies showing that binding of a single Ca^2+^ in the transport site I is sufficient to reverse SERCA inhibition by PLB (19). These Ca^2+^-induced structural rearrangements that follow relief of SERCA inhibition correspond to those associated with the formation of a competent transport site I and a vacant site II, which are required for binding of a second Ca^2+^ ion and subsequent activation of the pump (33,51,54,70).

In summary, we demonstrate that at saturating Ca^2+^ concentrations, binding of a single Ca^2+^ ion shifts the equilibrium toward a non-inhibited PLB-bound state of SERCA to induce reversal of SERCA. Our findings indicate that reversal this mechanism for reversal of SERCA-PLB inhibition solely dependents on Ca^2+^ concentration and is uncoupled from other regulatory mechanisms, such as the order-to-disorder structural transitions of PLB (12,39–43). The lack of a regulatory mechanism in saturating Ca^2+^-dependent relief of inhibition provides an explanation for the insufficient SERCA activation that contributes impaired Ca^2+^ transport (4,71,72) and maladaptations (73) characteristic of chronic heart failure.

## Experimental Procedures

### Setting up SERCA-PLB at super-physiological concentrations of Ca^2+^

We used an atomic model of full-length PLB bound to SERCA generated previously by our group (16) to simulate the inhibited SERCA–PLB complex at super-physiological Ca^2+^ conditions. We modeled transport site residues Glu309, Glu771 and Asp800 as unprotonated and residue Glu908 as protonated. In addition, we adjusted the p*K*a of other ionizable residues to a pH value of ~7.2 using PROPKA 3.1 (74, 75). The complex was inserted in a pre-equilibrated 120×120 Å bilayer of POPC lipids. We used the first layer phospholipids that surround SERCA in the E1 state (76) as a reference to insert the complex in the lipid bilayer. This initial system was solvated using TIP3P water molecules with a minimum margin of 15 Å between the protein and the edges of the periodic box in the z-axis. Ca^2+^ and Cl^−^ ions were added to produce a CaCl2 concentration of approximately 10 mM required to match the experimental conditions previously used to obtain crystal structures of SERCA at saturating Ca^2+^ conditions (24, 25). CHARMM36 force field topologies and parameters were used for the proteins (77), lipid (78), water, Ca^2+^, and Cl^−^.

### Setting up the SERCA-PLB complex at saturating Ca^2+^ conditions

We used the structure of the complex bound to a single Ca^2+^ ion obtained at super-physiological [Ca^2+^] as a starting structure to simulate the SERCA-PLB complex at saturating Ca^2+^ conditions. We found that a single Ca^2+^ ion bound to the transport sites of SERCA corresponds to a total Ca^2+^ concentration of ~400 μM. This total Ca^2+^ concentration is much higher than that estimated at rest (79), and is also in good agreement with previous estimates at elevated cytosolic Ca^2+^ (60–62). Residues Glu309, Glu771 and Asp800 were kept ionized, whereas residue Glu908 was modeled as neutral. The p*K*a of other ionizable residues were adjusted to a pH=7.2. The complex was inserted in a pre-equilibrated POPC lipid bilayer and solvated using TIP3P water molecules with a minimum margin of 15 Å between the protein and the edges of the periodic box in the z-axis. K+, and Cl^−^ ions were added to neutralize the system and to produce a KCl concentration of ~100 mM. CHARMM36 force field topologies and parameters were used for the protein (77), lipid (78), water, Ca^2+^, K+ and Cl^−^.

### Setting up the SERCA-PLB complex at free Ca^2+^ conditions

We used an atomic model of the full-length SERCA-PLB structure (16) to simulate the inhibited complex at free Ca^2+^ conditions. On the basis of our previous studies (16), we modeled transport site residues Glu309 and Asp800 as unprotonated and residues Glu771 and Glu908 as protonated. The lipid-water-protein complex was prepared using the same protocol as for the complex at saturating Ca^2+^ conditions. K^+^ and Cl^−^ ions were added to produce concentrations of 100 mM and 110 mM, respectively. CHARMM36 force field topologies and parameters were used for the proteins (77), lipid (78), water, K+, and Cl^−^.

### Molecular dynamics simulations

MD simulations of all systems were performed by using the program NAMD 2.12 (80), with periodic boundary conditions (81), particle mesh Ewald (82, 83), a non-bonded cutoff of 9 Å, and a 2 fs time step. The NPT ensemble was maintained with a Langevin thermostat (310K) and an anisotropic Langevin piston barostat (1 atm). Fully solvated systems were first subjected to energy minimization, followed by gradually warming up of the systems for 200 ps. This procedure was followed by 10 ns of equilibration with backbone atoms harmonically restrained using a force constant of 10 kcal mol^−1^ Å^−2^. We performed one 0.5-μs MD simulation of SERCA-PLB at 10 mM Ca^2+^, and 12 independent 1-μs MD simulations: six of Ca^2+^-bound SERCA-PLB and six of SERCA-PLB in the absence of Ca^2+^.

### Structural analysis and visualization

VMD (84) was used for analysis, visualization, and rendering of the structures. To visualize the Ca^2+^-protein and Ca^2+^-lipid interactions, we created a map of the weighted mass density of Ca^2+^ using a grid resolution of 1 Å and a cutoff distance of 3.5 Å between Ca^2+^ and the protein/lipid atoms. This is achieved by replacing each atom in the selection with a normalized Gaussian distribution of width equal to the atomic radius. The distributions are then additively distributed on a grid. The final map is calculated by computing the mass density of Ca^2+^ for each step in the trajectory and averaged over the entire simulation time.

We calculated the fraction of native inhibitory contacts (*Qinh*) between PLB residues Leu31, Asn34, Phe35 and Ile38 and SERCA to measure the effect of calcium binding on the stability of the SERCA-PLB interface. *Q_inh_* is defined by a list of native contact pairs (*i*,*j*) in the crystal structure of the complex. All pairs of heavy atoms *i* and *j* belonging to residues X*i* and X*j* are in contact if the distance between *i* and *j* is <0.7 nm. *Q_inh_* is expressed as a number between 1 and 0, and it is calculated as the total number of native contacts for a given time frame divided by the total number of contacts in the crystal structure of the complex [PDB code: 4kyt (15)].

## Acknowledgments

We thank Joseph M. Autry and Yoshiki Kabashima for helpful discussions. Computational resources were provided by the Minnesota Supercomputing Institute.

## Funding Sources and Disclaimers

This study was funded by the National Institutes of Health grant R01GM120142 to L.M.E.-F. The content is solely the responsibility of the authors and does not necessarily represent the official views of the National Institutes of Health.

## Footnotes

This study was funded by National Institutes of Health grant R01GM120142 to LMEF. The content is solely the responsibility of the authors and does not necessarily represent the official views of the National Institutes of Health.

Abbreviations used are: SERCA, sarcoplasmic reticulum Ca^2+^-ATPase; PLB, phospholamban; MD, molecular dynamics; SLN, sarcolipin; SR, sarcoplasmic reticulum; TM, transmembrane; POPC, palmitoyl-2-oleoyl-sn-glycerol-phosphatidylcholine; root-mean square deviation, RMSD; root-mean square fluctuation, RMSF.

## References

1. Moller, J. V., Olesen, C., Winther, A. M., and Nissen, P. (2010) The sarcoplasmic Ca2+-ATPase: design of a perfect chemi-osmotic pump. Q Rev Biophys 43, 501–566

2. Cantilina, T., Sagara, Y., Inesi, G., and Jones, L. R. (1993) Comparative studies of cardiac and skeletal sarcoplasmic reticulum ATPases. Effect of a phospholamban antibody on enzyme activation by Ca2+. J Biol Chem 268, 17018–17025.

3. Sasaki, T., Inui, M., Kimura, Y., Kuzuya, T., and Tada, M. (1992) Molecular mechanism of regulation of Ca2+ pump ATPase by phospholamban in cardiac sarcoplasmic reticulum. Effects of synthetic phospholamban peptides on Ca2+ pump ATPase. The Journal of biological chemistry 267, 1674–1679

4. MacLennan, D. H., and Kranias, E. G. (2003) Phospholamban: a crucial regulator of cardiac contractility. Nature reviews. Molecular cell biology 4, 566–577

5. Akin, B. L., and Jones, L. R. (2012) Characterizing phospholamban to sarco(endo)plasmic reticulum Ca2+-ATPase 2a (SERCA2a) protein binding interactions in human cardiac sarcoplasmic reticulum vesicles using chemical cross-linking. J Biol Chem 287, 7582–7593

6. Chen, Z., Akin, B. L., Stokes, D. L., and Jones, L. R. (2006) Cross-linking of C-terminal residues of phospholamban to the Ca2+ pump of cardiac sarcoplasmic reticulum to probe spatial and functional interactions within the transmembrane domain. J Biol Chem 281, 14163–14172

7. Toyoshima, C., Asahi, M., Sugita, Y., Khanna, R., Tsuda, T., and MacLennan, D. H. (2003) Modeling of the inhibitory interaction of phospholamban with the Ca2+ ATPase. Proc Natl Acad Sci U S A 100, 467–472

8. Zamoon, J., Nitu, F., Karim, C., Thomas, D. D., and Veglia, G. (2005) Mapping the interaction surface of a membrane protein: unveiling the conformational switch of phospholamban in calcium pump regulation. Proc Natl Acad Sci U S A 102, 4747–4752

9. Asahi, M., Kimura, Y., Kurzydlowski, K., Tada, M., and MacLennan, D. H. (1999) Transmembrane helix M6 in sarco(endo)plasmic reticulum Ca(2+)-ATPase forms a functional interaction site with phospholamban. Evidence for physical interactions at other sites. The Journal of biological chemistry 274, 32855–32862

10. Morita, T., Hussain, D., Asahi, M., Tsuda, T., Kurzydlowski, K., Toyoshima, C., and Maclennan, D. H. (2008) Interaction sites among phospholamban, sarcolipin, and the sarco(endo)plasmic reticulum Ca(2+)-ATPase. Biochemical and biophysical research communications 369, 188–194

11. Dong, X., and Thomas, D. D. (2014) Time-resolved FRET reveals the structural mechanism of SERCA-PLB regulation. Biochemical and biophysical research communications 449, 196–201

12. Gustavsson, M., Verardi, R., Mullen, D. G., Mote, K. R., Traaseth, N. J., Gopinath, T., and Veglia, G. (2013) Allosteric regulation of SERCA by phosphorylation-mediated conformational shift of phospholamban. Proc Natl Acad Sci U S A 110, 17338–17343

13. Ha, K. N., Traaseth, N. J., Verardi, R., Zamoon, J., Cembran, A., Karim, C. B., Thomas, D. D., and Veglia, G. (2007) Controlling the inhibition of the sarcoplasmic Ca2+-ATPase by tuning phospholamban structural dynamics. J Biol Chem 282, 37205–37214

14. Karim, C. B., Kirby, T. L., Zhang, Z., Nesmelov, Y., and Thomas, D. D. (2004) Phospholamban structural dynamics in lipid bilayers probed by a spin label rigidly coupled to the peptide backbone. Proc Natl Acad Sci U S A 101, 14437–14442

15. Akin, B. L., Hurley, T. D., Chen, Z., and Jones, L. R. (2013) The structural basis for phospholamban inhibition of the calcium pump in sarcoplasmic reticulum. The Journal of biological chemistry 288, 30181–30191

16. Espinoza-Fonseca, L. M., Autry, J. M., Ramirez-Salinas, G. L., and Thomas, D. D. (2015) Atomic-level mechanisms for phospholamban regulation of the calcium pump. Biophysical journal 108, 1697–1708

17. Espinoza-Fonseca, L. M., Autry, J. M., and Thomas, D. D. (2015) Sarcolipin and phospholamban inhibit the calcium pump by populating a similar metal ion-free intermediate state. Biochemical and biophysical research communications 463, 37–41

18. Asahi, M., McKenna, E., Kurzydlowski, K., Tada, M., and MacLennan, D. H. (2000) Physical interactions between phospholamban and sarco(endo)plasmic reticulum Ca2+-ATPases are dissociated by elevated Ca2+, but not by phospholamban phosphorylation, vanadate, or thapsigargin, and are enhanced by ATP. The Journal of biological chemistry 275, 15034–15038

19. Chen, Z., Akin, B. L., and Jones, L. R. (2010) Ca2+ binding to site I of the cardiac Ca2+ pump is sufficient to dissociate phospholamban. J Biol Chem 285, 3253–3260

20. Bidwell, P., Blackwell, D. J., Hou, Z., Zima, A. V., and Robia, S. L. (2011) Phospholamban binds with differential affinity to calcium pump conformers. The Journal of biological chemistry 286, 35044–35050

21. James, Z. M., McCaffrey, J. E., Torgersen, K. D., Karim, C. B., and Thomas, D. D. (2012) Protein-Protein Interactions in Calcium Transport Regulation Probed by Saturation Transfer Electron Paramagnetic Resonance. Biophysical journal 103, 1370–1378

22. Mueller, B., Karim, C. B., Negrashov, I. V., Kutchai, H., and Thomas, D. D. (2004) Direct detection of phospholamban and sarcoplasmic reticulum Ca-ATPase interaction in membranes using fluorescence resonance energy transfer. Biochemistry 43, 8754–8765

23. Kirschenlohr, H. L., Grace, A. A., Vandenberg, J. I., Metcalfe, J. C., and Smith, G. A. (2000) Estimation of systolic and diastolic free intracellular Ca2+ by titration of Ca2+ buffering in the ferret heart. The Biochemical journal 346 Pt 2, 385–391

24. Toyoshima, C., and Mizutani, T. (2004) Crystal structure of the calcium pump with a bound ATP analogue. Nature 430, 529–535

25. Toyoshima, C., Nakasako, M., Nomura, H., and Ogawa, H. (2000) Crystal structure of the calcium pump of sarcoplasmic reticulum at 2.6 A resolution. Nature 405, 647–655

26. Sorensen, T. L., Clausen, J. D., Jensen, A. M., Vilsen, B., Moller, J. V., Andersen, J. P., and Nissen, P. (2004) Localization of a K+ -binding site involved in dephosphorylation of the sarcoplasmic reticulum Ca2+ -ATPase. J Biol Chem 279, 46355–46358

27. Smolin, N., and Robia, S. L. (2015) A structural mechanism for calcium transporter headpiece closure. J Phys Chem B 119, 1407–1415

28. Deng, H., Chen, G., Yang, W., and Yang, J. J. (2006) Predicting calcium-binding sites in proteins - a graph theory and geometry approach. Proteins 64, 34–42

29. Dudev, T., Lin, Y. L., Dudev, M., and Lim, C. (2003) First-second shell interactions in metal binding sites in proteins: a PDB survey and DFT/CDM calculations. J Am Chem Soc 125, 3168–3180

30. Nayal, M., and Di Cera, E. (1994) Predicting Ca(2+)-binding sites in proteins. Proc Natl Acad Sci US A 91, 817–821

31. Carugo, O., Djinovic, K., and Rizzi, M. (1993) Comparison of the Coordinative Behavior of Calcium(Ii) and Magnesium(Ii) from Crystallographic Data. J Chem Soc Dalton, 2127–2135

32. Katz, A. K., Glusker, J. P., Beebe, S. A., and Bock, C. W. (1996) Calcium ion coordination: A comparison with that of beryllium, magnesium, and zinc. J Am Chem Soc 118, 5752–5763

33. Toyoshima, C., Nakasako, M., Nomura, H., and Ogawa, H. (2000) Crystal structure of the calcium pump of sarcoplasmic reticulum at 2.6 angstrom resolution. Nature 405, 647–655

34. Kimura, Y., Kurzydlowski, K., Tada, M., and MacLennan, D. H. (1996) Phospholamban regulates the Ca2+-ATPase through intramembrane interactions. Journal of Biological Chemistry 271, 21726–21731

35. Strock, C., Cavagna, M., Peiffer, W. E., Sumbilla, C., Lewis, D., and Inesi, G. (1998) Direct demonstration of Ca2+ binding defects in sarco-endoplasmic reticulum Ca2+ ATPase mutants overexpressed in COS-1 cells transfected with adenovirus vectors. J Biol Chem 273, 15104–15109

36. Pal, T. K., and Sankararamakrishnan, R. (2008) Self-contacts in Asx and Glx residues of high-resolution protein structures: Role of local environment and tertiary interactions. J Mol Graph Model 27, 20–33

37. Traaseth, N. J., Shi, L., Verardi, R., Mullen, D. G., Barany, G., and Veglia, G. (2009) Structure and topology of monomeric phospholamban in lipid membranes determined by a hybrid solution and solid-state NMR approach. Proc Natl Acad Sci U S A 106, 10165–10170

38. Zamoon, J., Mascioni, A., Thomas, D. D., and Veglia, G. (2003) NMR solution structure and topological orientation of monomeric phospholamban in dodecylphosphocholine micelles. Biophys J 85, 2589–2598

39. Karim, C. B., Zhang, Z., Howard, E. C., Torgersen, K. D., and Thomas, D. D. (2006) Phosphorylation-dependent conformational switch in spin-labeled phospholamban bound to SERCA. J Mol Biol 358, 1032–1040

40. De Simone, A., Gustavsson, M., Montalvao, R. W., Shi, L., Veglia, G., and Vendruscolo, M. (2013) Structures of the excited states of phospholamban and shifts in their populations upon phosphorylation. Biochemistry 52, 6684–6694

41. Metcalfe, E. E., Traaseth, N. J., and Veglia, G. (2005) Serine 16 phosphorylation induces an order-to-disorder transition in monomeric phospholamban. Biochemistry 44, 4386–4396

42. Traaseth, N. J., Thomas, D. D., and Veglia, G. (2006) Effects of Ser16 phosphorylation on the allosteric transitions of phospholamban/Ca(2+)-ATPase complex. J Mol Biol 358, 1041–1050

43. Paterlini, M. G., and Thomas, D. D. (2005) The alpha-helical propensity of the cytoplasmic domain of phospholamban: a molecular dynamics simulation of the effect of phosphorylation and mutation. Biophys J 88, 3243–3251

44. Lee, A. G. (2004) How lipids affect the activities of integral membrane proteins. Bba-Biomembranes 1666, 62–87

45. Kimura, Y., Kurzydlowski, K., Tada, M., and MacLennan, D. H. (1997) Phospholamban inhibitory function is activated by depolymerization. Journal of Biological Chemistry 272, 15061–15064

46. Bublitz, M., Musgaard, M., Poulsen, H., Thogersen, L., Olesen, C., Schiott, B., Morth, J. P., Moller, J. V., and Nissen, P. (2013) Ion pathways in the sarcoplasmic reticulum Ca2+-ATPase. The Journal of biological chemistry 288, 10759–10765

47. Einholm, A. P., Vilsen, B., and Andersen, J. P. (2004) Importance of transmembrane segment M1 of the sarcoplasmic reticulum Ca2+-ATPase in Ca2+ occlusion and phosphoenzyme processing. J Biol Chem 279, 15888–15896

48. Kekenes-Huskey, P. M., Metzger, V. T., Grant, B. J., and Andrew McCammon, J. (2012) Calcium binding and allosteric signaling mechanisms for the sarcoplasmic reticulum Ca(2)+ ATPase. Protein Sci 21, 1429–1443

49. Obara, K., Miyashita, N., Xu, C., Toyoshima, I., Sugita, Y., Inesi, G., and Toyoshima, C. (2005) Structural role of countertransport revealed in Ca(2+) pump crystal structure in the absence of Ca(2+). Proc Natl Acad Sci U S A 102, 14489–14496

50. Toyoshima, C., and Cornelius, F. (2013) New crystal structures of PII-type ATPases: excitement continues. Current opinion in structural biology 23, 507–514

51. Espinoza-Fonseca, L. M., Autry, J. M., and Thomas, D. D. (2014) Microsecond molecular dynamics simulations of Mg(2)(+)- and K(+)-bound E1 intermediate states of the calcium pump. PloS one 9, e95979

52. Inesi, G., Lewis, D., Toyoshima, C., Hirata, A., and de Meis, L. (2008) Conformational fluctuations of the Ca2+-ATPase in the native membrane environment. Effects of pH, temperature, catalytic substrates, and thapsigargin. The Journal of biological chemistry 283, 1189–1196

53. Fernandez-De Gortari, E., and Espinoza-Fonseca, L. M. (2017) Preexisting domain motions underlie protonation-dependent structural transitions of the P-type Ca2+-ATPase. Phys Chem Chem Phys 19, 10153–10162

54. Espinoza-Fonseca, L. M., and Thomas, D. D. (2011) Atomic-level characterization of the activation mechanism of SERCA by calcium. PLoS One 6, e26936

55. Winther, A. M., Bublitz, M., Karlsen, J. L., Moller, J. V., Hansen, J. B., Nissen, P., and Buch-Pedersen, M. J. (2013) The sarcolipin-bound calcium pump stabilizes calcium sites exposed to the cytoplasm. Nature 495, 265–269

56. Toyoshima, C., Iwasawa, S., Ogawa, H., Hirata, A., Tsueda, J., and Inesi, G. (2013) Crystal structures of the calcium pump and sarcolipin in the Mg2+-bound E1 state. Nature 495, 260–264

57. Inesi, G., Kurzmack, M., Coan, C., and Lewis, D. E. (1980) Cooperative calcium binding and ATPase activation in sarcoplasmic reticulum vesicles. J Biol Chem 255, 3025–3031

58. Mintz, E., and Guillain, F. (1997) Ca2+ transport by the sarcoplasmic reticulum ATPase. Bba-Bioenergetics 1318, 52–70

59. Inesi, G., Sumbilla, C., and Kirtley, M. E. (1990) Relationships of Molecular-Structure and Function in Ca-2+-Transport Atpase. Physiol Rev 70, 749–760

60. Hove-Madsen, L., and Bers, D. M. (1993) Passive Ca buffering and SR Ca uptake in permeabilized rabbit ventricular myocytes. Am J Physiol 264, C677–686

61. Balke, C. W., Egan, T. M., and Wier, W. G. (1994) Processes That Remove Calcium from the Cytoplasm during Excitation-Contraction Coupling in Intact Rat-Heart Cells. J Physiol-London 474, 447–462

62. Wendtgallitelli, M. F., and Isenberg, G. (1991) Total and Free Myoplasmic Calcium during a Contraction Cycle - X-Ray-Microanalysis in Guinea-Pig Ventricular Myocytes. J Physiol-London 435, 349–372

63. Sugita, Y., Ikeguchi, M., and Toyoshima, C. (2010) Relationship between Ca2+-affinity and shielding of bulk water in the Ca2+-pump from molecular dynamics simulations. P Natl Acad Sci USA 107, 21465–21469

64. Chen, Z., Akin, B. L., and Jones, L. R. (2007) Mechanism of reversal of phospholamban inhibition of the cardiac Ca2+-ATPase by protein kinase a and by anti-phospholamban monoclonal antibody 2D12. Journal of Biological Chemistry 282, 20968–20976

65. Gorski, P. A., Glaves, J. P., Vangheluwe, P., and Young, H. S. (2013) Sarco(endo)plasmic Reticulum Calcium ATPase (SERCA) Inhibition by Sarcolipin Is Encoded in Its Luminal Tail. Journal of Biological Chemistry 288, 8456–8467

66. Marcus, Y. (1988) Ionic-Radii in Aqueous-Solutions. Chem Rev 88, 1475–1498

67. Dudev, T., and Lim, C. (1999) Incremental binding free energies in Mg2+ complexes: A DFT study. J Phys Chem A 103, 8093–8100

68. Dudev, T., and Lim, C. (2004) Monodentate versus bidentate carboxylate binding in magnesium and calcium proteins: What are the basic principles? Journal of Physical Chemistry B 108, 4546–4557

69. Inesi, G., Ma, H., Lewis, D., and Xu, C. (2004) Ca2+ occlusion and gating function of Glu309 in the ADP-fluoroaluminate analog of the Ca2+-ATPase phosphoenzyme intermediate. J Biol Chem 279, 31629–31637

70. Sugita, Y., Ikeguchi, M., and Toyoshima, C. (2010) Relationship between Ca2+-affinity and shielding of bulk water in the Ca2+-pump from molecular dynamics simulations. Proc Natl Acad Sci U S A 107, 21465–21469

71. Periasamy, M., and Huke, S. (2001) SERCA pump level is a critical determinant of Ca2+ homeostasis and cardiac contractility. J Mol Cell Cardiol 33, 1053–1063

72. Sande, J. B., Sjaastad, I., Hoen, I. B., Bokenes, J., Tonnessen, T., Holt, E., Lunde, P. K., and Christensen, G. (2002) Reduced level of serine(16) phosphorylated phospholamban in the failing rat myocardium: a major contributor to reduced SERCA2 activity. Cardiovasc Res 53, 382–391

73. Sabbah, H. N., Gupta, R. C., Kohli, S., Wang, M., Zhang, K., and Rastogi, S. (2014) Heart rate reduction with ivabradine improves left ventricular function and reverses multiple pathological maladaptations in dogs with chronic heart failure. ESC Heart Fail 0, 94–102

74. Olsson, M. H. M., Sondergaard, C. R., Rostkowski, M., and Jensen, J. H. (2011) PROPKA3: Consistent Treatment of Internal and Surface Residues in Empirical pK(a) Predictions. J Chem Theory Comput 7, 525–537

75. Sondergaard, C. R., Olsson, M. H. M., Rostkowski, M., and Jensen, J. H. (2011) Improved Treatment of Ligands and Coupling Effects in Empirical Calculation and Rationalization of pK(a) Values. J Chem Theory Comput 7, 2284–2295

76. Norimatsu, Y., Hasegawa, K., Shimizu, N., and Toyoshima, C. (2017) Protein-phospholipid interplay revealed with crystals of a calcium pump. Nature 545, 193–198

77. Best, R. B., Zhu, X., Shim, J., Lopes, P. E., Mittal, J., Feig, M., and Mackerell, A. D., Jr. (2012) Optimization of the additive CHARMM all-atom protein force field targeting improved sampling of the backbone phi, psi and side-chain chi(1) and chi(2) dihedral angles. J Chem Theory Comput 8, 3257–3273

78. Klauda, J. B., Venable, R. M., Freites, J. A., O’Connor, J. W., Tobias, D. J., Mondragon-Ramirez, C., Vorobyov, I., MacKerell, A. D., Jr., and Pastor, R. W. (2010) Update of the CHARMM all-atom additive force field for lipids: validation on six lipid types. J Phys Chem B 114, 7830–7843

79. Bers, D. M. (2002) Cardiac excitation-contraction coupling. Nature 415, 198–205

80. Phillips, J. C., Braun, R., Wang, W., Gumbart, J., Tajkhorshid, E., Villa, E., Chipot, C., Skeel, R. D., Kale, L., and Schulten, K. (2005) Scalable molecular dynamics with NAMD. J Comput Chem 26, 1781–1802

81. Weber, W., Hünenberger, P. H., and McCammon, J. A. (2000) Molecular Dynamics Simulations of a Polyalanine Octapeptide under Ewald Boundary Conditions: Influence of Artificial Periodicity on Peptide Conformation. J Phys Chem B 104, 3668–3675

82. Darden, T., York, D., and Pedersen, L. (1993) Particle mesh Ewald: An N.log(N) method for Ewald sums in large systems. J Chem Phys 98, 10089–10092

83. Essmann, U., Perera, L., and Berkowitz, M. L. (1995) A smooth particle mesh Ewald method. J Chem Phys 103, 8577–8593

84. Humphrey, W., Dalke, A., and Schulten, K. (1996) VMD: visual molecular dynamics. J Mol Graph 14, 33–38, 27–38

